# Enhancing biological signals and detection rates in single-cell RNA-seq experiments with cDNA library equalization

**DOI:** 10.1101/2020.10.05.326553

**Authors:** Rhonda Bacher, Li-Fang Chu, Cara Argus, Jennifer M. Bolin, Parker Knight, James A. Thomson, Ron Stewart, Christina Kendziorski

**Affiliations:** Department of Biostatistics, University of Florida, FL, USA; Morgridge Institute for Research, Madison, WI, USA; Department of Mathematics, University of Florida, FL, USA; Department of Biostatistics, University of Wisconsin-Madison, WI, USA

## Abstract

Considerable effort has been devoted to refining experimental protocols having reduced levels of technical variability and artifacts in single-cell RNA-sequencing data (scRNA-seq). We here present evidence that equalizing the concentration of cDNA libraries prior to pooling, a step not consistently performed in single-cell experiments, improves gene detection rates, enhances biological signals, and reduces technical artifacts in scRNA-seq data. To evaluate the effect of equalization on various protocols, we developed Scaffold, a simulation framework that models each step of an scRNA-seq experiment. Numerical experiments demonstrate that equalization reduces variation in sequencing depth and gene-specific expression variability. We then performed a set of experiments in vitro with and without the equalization step and found that equalization increases the number of genes that are detected in every cell by 17-31%, improves discovery of biologically relevant genes, and reduces nuisance signals associated with cell cycle. Further support is provided in an analysis of publicly available data.

## Introduction

Single-cell RNA-sequencing (scRNA-seq) protocols have evolved rapidly over the last ten years, with increased throughput and sensitivity allowing for unprecedented insights into cell type heterogeneity across tissues(1). In spite of the advances, substantial technical variability and biases remain, which present challenges in data analysis and can obscure biological signals(2–5). From mRNA capture, reverse transcription, and PCR amplification, to additional single-cell library preparation and multiplex sequencing, there are numerous opportunities for technical noise to arise in scRNA-seq experiments. Inefficiencies or biases at any of the steps in the protocol may lead to increased technical artifacts and noise affecting expression variability and increasing the number of zeros(6,7).

Numerous computational approaches including data smoothing and imputation have been developed to address excess variability and zeros in scRNA-seq data(8,9). However, they do so with the risk of introducing or perpetuating bias(10), thus making it preferable to optimize experimental protocols when feasible. A few studies have evaluated the downstream effects of various amplification techniques(11) or reverse transcriptases(12) on scRNA-seq data. However, to our knowledge no study has assessed the effect of equalizing cDNA concentrations in single-cell protocols. In bulk RNA-seq experiments, equalization of cDNA concentrations across libraries is a standard procedure that has been shown to reduce sequencing coverage variability and increase transcriptome diversity(13–15) by providing more even sequencing coverage of all samples. Equalization also leads to decreased sequencing of highly abundant transcripts and increases the efficiency at which low and moderately expressed genes are sequenced in bulk experiments(14).

For single-cell RNA-seq we hypothesized that equalization may improve sensitivity by increasing gene detection and thus began our investigation into the technical artifacts in scRNA-seq data by developing a simulation framework, Scaffold, that generates counts by modelling each step of the experimental protocol. Simulation frameworks offer a significant advantage to studying sources of variability compared to experimental approaches as they allow an investigator to quickly assess a large number of scenarios at considerably low cost. While a number of good methods are available for simulating scRNA-seq data(16–18), most do not model each step in the experimental protocol, and therefore are not useful for assessing how each step of the process affects the final counts. Two frameworks have attempted to study the data generation process but are limited in scope, either relying on spike-ins (19) or combining all sources of variation into a single parameter(20). Scaffold models each step in an scRNA-seq data generating process by representing each step of the protocol mathematically, from the initial cell-to-cell heterogeneity to the final sequencing (Methods). We focus on the SMART-SEQ(21) protocol as it uses oligo-dT priming and template switching as the backbone chemistry to generate cDNA from single cells which is used in multiple major scRNA-seq platforms, including Fluidigm C1 and 10X Chromium.

Based on our simulation results which suggest that equalization is a critical step in the scRNA-seq protocol, we designed a set of scRNA-seq experiments in which we varied the extent at which cDNA libraries were equalized. The experiments demonstrate that equalization results in more consistent detection of genes, reduced expression variability, and reduced variability in the count-depth rate(3), the relationship between a gene’s observed expression and sequencing depth. Finally, we confirm the effect of equalization in a survey of publicly available scRNA-seq datasets.

## Methods

### EC and TB cell experiments

We focused on a subset of 96 single cells, from hESC-derived endothelial cells (EC) or trophoblast-like cells (TB) generated using the Fluidigm C1 system. The original data is considered to be unequalized (unEQ), where the single-cell cDNA libraries were first diluted to a range of 0.125–0.375 ng for subsequent library preparation protocols. The unEQ data was published in a previous study (GEO: GSE75748)(22). In the subsequent EQ experiments performed here, including EQ, EQ-Vary and EQ-75%, we retrieved the harvested cDNA, which are amplified full-length single-cell cDNAs identical to those used for the unEQ experiments (Supplementary Figure 7), but further diluted and adjusted so only 0.1 ng of cDNA were used as input across all the cells for subsequent library preparation protocols. In all the experiments, 1.25 µL of indicated input cDNA were used in a 5.0 µL Tagmentation reaction (Nextera XT DNA Sample Preparation Kit, Illumina) followed with a 12.5 µL dual-indexing PCR amplification reaction (Nextera XT DNA Sample Preparation Index Kit, Illumina). In the unEQ, EQ and EQ-75% experiments, 2.0 µL of the amplified/tagmented cDNA were used for pooling. In the EQ-Vary experiment, a single scaling factor was applied to generate variable amounts of the pooling volume. These pooled single-cell libraries were used in an AMpure XP Bead-based Dual Bead Cleanup and Size Selection reaction (Agencourt AMPure XP PCR Purification modified Instructions for Use, Beckman Coulter). In both bead cleanup reactions, 90% of AMPure XP beads were added to the amplified single-cell libraries to select for an approximate size range of 150-700 bp and incubated for 15 minutes at room temperature. Libraries bound to beads were then placed on a magnet for 5 minutes, washed twice with 70% Ethanol, eluted with Suspension Buffer (Nextera XT DNA Sample Preparation Index Kit, Illumina), and transferred to a new tube. Final amplified and pooled single-cell libraries were quantified with the Qubit dsDNA HS Assay Kit (Q32854, Thermofisher) and Bioanalyzer High Sensitivity DNA Analysis Kit (5067-4626, Agilent). The unEQ was multiplexed with 18-20 samples per lane and sequenced on an Illumina HiSeq2500 with single-end 51 bp reads while the EQ, EQ-75%, and EQ-Vary were all pooled with 96 samples per lane and sequenced on an Illumina HiSeq3000 with paired-end 65 or 78 bp reads.

Reads were mapped against the GRCh38 Ensembl reference of protein-coding genes via Bowtie 1.2.3(34), allowing up to two mismatches. The expected counts were estimated via RSEM 1.2.31(35). To control for any difference due to differing read lengths, all reads were first trimmed to have a length of 51bp. In the initial unequalized experiment, cells that had less than 5,000 genes with TPM >1 or that upon inspection of cell images displayed doublets or appeared dead were removed in quality control.

### Quality control on cells across equalization experiments

Using the scater v1.12.2 R package(36) we removed cells from any experiments in which the log10 sequencing depth was < 5.4 or the percent of counts in the top 50 genes was > 31%, the thresholds corresponding to being two standard deviations away from the median (Supplementary Figure 8). The expected counts in all experiments were rounded to the nearest whole number for all subsequent analyses.

### Comparison of cell-specific and gene-specific detection rates

The cell-specific detection rate was calculated as the proportion of genes with nonzero expression within each cell. Similarly, the gene-specific detection rate was calculated as the proportion of cells with nonzero expression for each gene. When comparing the differences in gene-specific detection rates between any two datasets, we accounted for differences in the sequencing depth by using the largest subset of cells for which an equal number of cells had an increase or decrease in sequencing depth.

### Analysis of highly variable genes

For the analysis of highly variable genes, gene expression estimates were first normalized using SCnorm v1.6.0(3). We then fit a mean-dependent trend across all genes mean-variance relationship. The trend represents technical variability and a gene’s biological variability was calculated as the difference between its total variance to the technical fitted trend. This was done using the scran package v1.12.1 in R using the functions trendVar and decomposeVar(37). Genes were considered highly variable in any dataset if they had an FDR < .10. In order to compare genes variability across datasets, we ranked a gene’s relative variability to all other genes in the dataset and calculated the difference in the two ranks.

### Estimating the count-depth rate

The gene-specific count-depth rate was estimated within EC and TB separately using a median quantile regression on the log nonzero gene expression versus log sequencing depth using the getSlopes function in the SCnorm v1.6.0 R package. For each condition, we filtered out genes that had less than 10 nonzero expression counts across all cells and genes with median nonzero expression less than two. Visualization of the count-depth rate distributions is shown using smoothed density plots of the slopes within gene groups, where genes were split into 10 equally sized groups based on their nonzero median expression. The variability of the count-depth rate is quantified using the median absolute deviation statistic (MAD). First, the mode of the slope distribution was estimated for each gene group, then the MAD was calculated as the median of the absolute differences between the slope modes and one, where one is the expected value of the count-depth rate. All density plots of the slope distribution are done with smoothing parameters adjust = 1, and estimated over the grid (−3,3) using the density function in R. All analyses were carried out using R version 3.6.3.

### Analysis of publicly available datasets

For each dataset, cells with less than 10,000 total counts were removed and counts were rounded to the nearest whole number. For estimating the count-depth rate, again we filtered out genes that had less than 10 nonzero expression counts across all cells and genes with median nonzero expression less than two. In Figure 4, the representative datasets displayed from each study are: EF cells from Islam, Earlyblast-Embryo2 in Deng, M11W-Embryo2 in Guo, Unstim-Rep1 in Shalek, and TB2 in Chu. The Picelli and H1-Bulk each only had one dataset in the study. The comparison of properties in Table 1 for the equalized versus unequalized datasets in publicly available studies was done using a two-sided t-test.

**Table 1.**
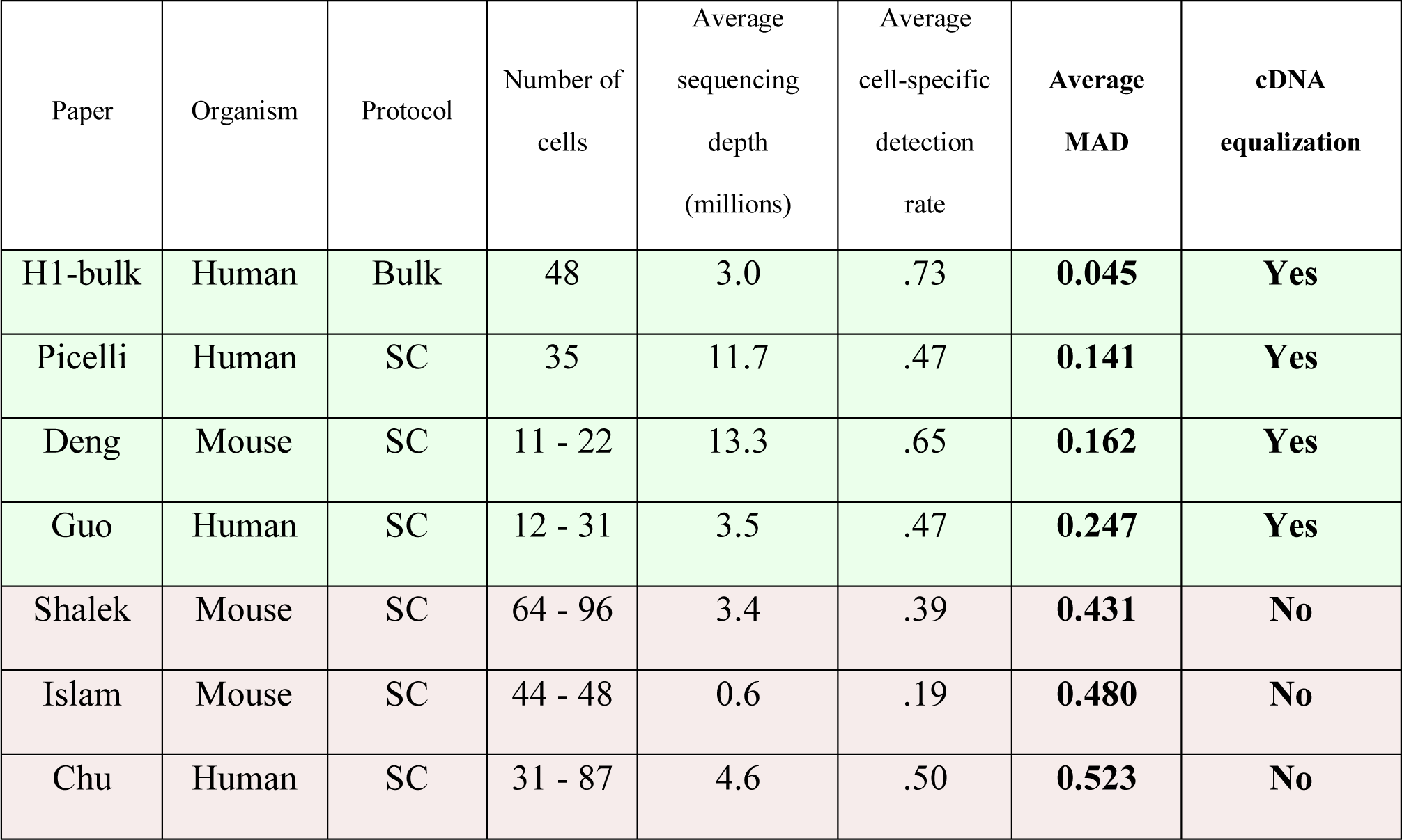
Summary of publicly available datasets. The first column contains the reference study. Column 2 shows the organism. Column 3 shows the sequencing protocol used. Column 4 shows the number of cells per dataset included in the study. Column 5 is average sequencing depth across all cells. Column 6 is the average cell-specific detection rate across all cells. Column 7 is the average MAD and Column 8 indicates whether cDNA equalization was performed.

| Paper | Organism | Protocol | Number of cells | Average sequencing depth (millions) | Average cell-specific detection rate | Average MAD | cDNA equalization |
| --- | --- | --- | --- | --- | --- | --- | --- |
| H1-bulk | Human | Bulk | 48 | 3.0 | .73 | **0.045** | **Yes** |
| Picelli | Human | SC | 35 | 11.7 | .47 | **0.141** | **Yes** |
| Deng | Mouse | SC | 11 - 22 | 13.3 | .65 | **0.162** | **Yes** |
| Guo | Human | SC | 12 - 31 | 3.5 | .47 | **0.247** | **Yes** |
| Shalek | Mouse | SC | 64 - 96 | 3.4 | .39 | **0.431** | **No** |
| Islam | Mouse | SC | 44 - 48 | 0.6 | .19 | **0.480** | **No** |
| Chu | Human | SC | 31 - 87 | 4.6 | .50 | **0.523** | **No** |

**Figure 1.**
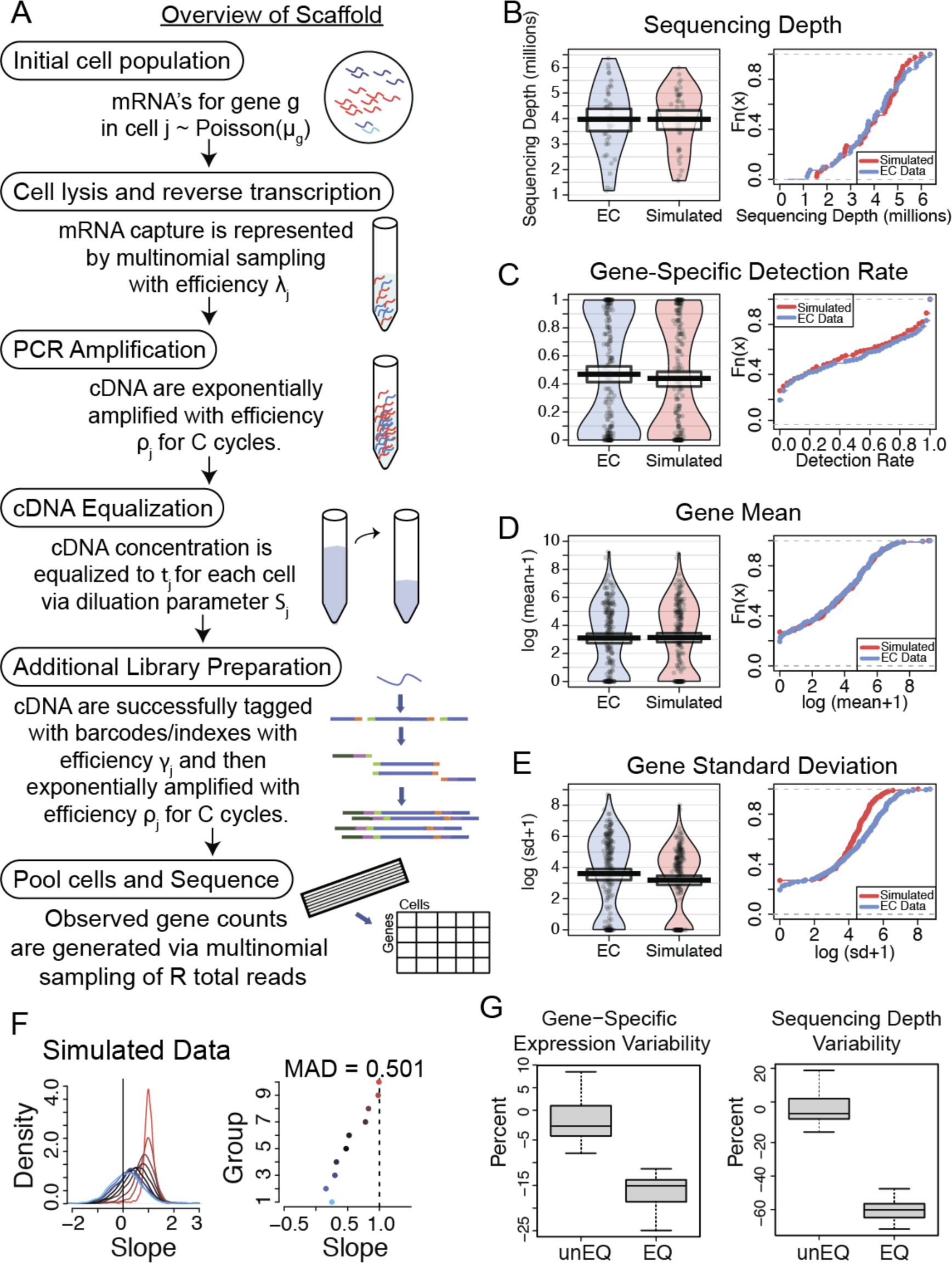
A. Overview of the Scaffold simulation framework. Further details are provided in Methods. B-E. Cell-specific and gene-specific properties of the data simulated based on the unEQ EC dataset. F. Density plots of the distribution of estimated count-depth rates (quantified as the gene-specific slope of a median quantile regression) for the unEQ EC dataset for genes grouped by expression level (left) and the mode of each group’s slope distribution (right). The median absolute deviation of the slope modes from one (MAD) is used to quantify the variability in the count-depth rate. G. The percent change in gene-specific variability (left) and sequencing depth (right) is shown for pairs of equalized and unequalized datasets. Pairs of unequalized experiments were also simulated and compared to demonstrate the percent of change due to random sampling.

**Figure 2.**
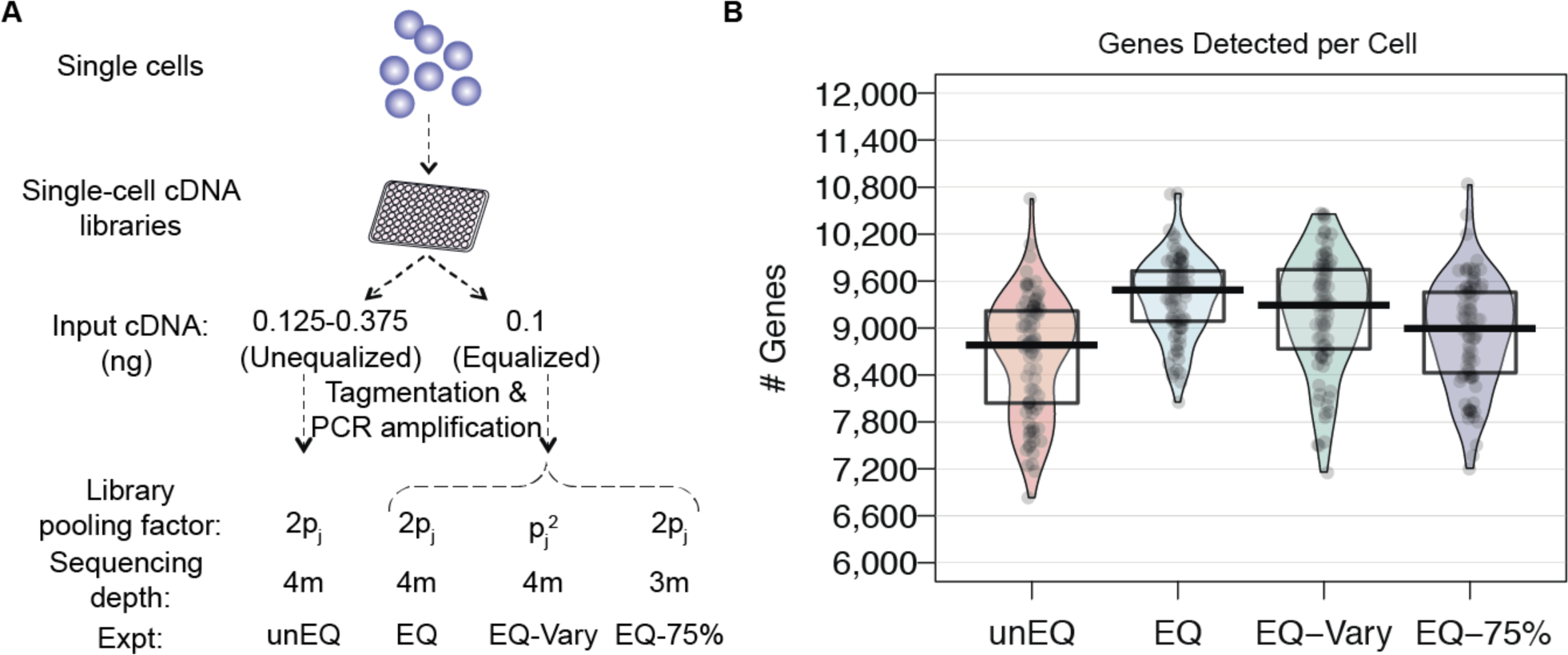
Overview of experiment to assess the effect of cDNA equalization and comparisons of cell-level detection rates. A.) Four experiments were conducted involving cells from two different conditions (EC and TB). Using the same initial pools of single-cell cDNA, we created unequalized and equalized sequencing libraries. B.) Violin plots with points overlaid of the number of genes detected per cell for all cells in each experiment.

**Figure 3.**
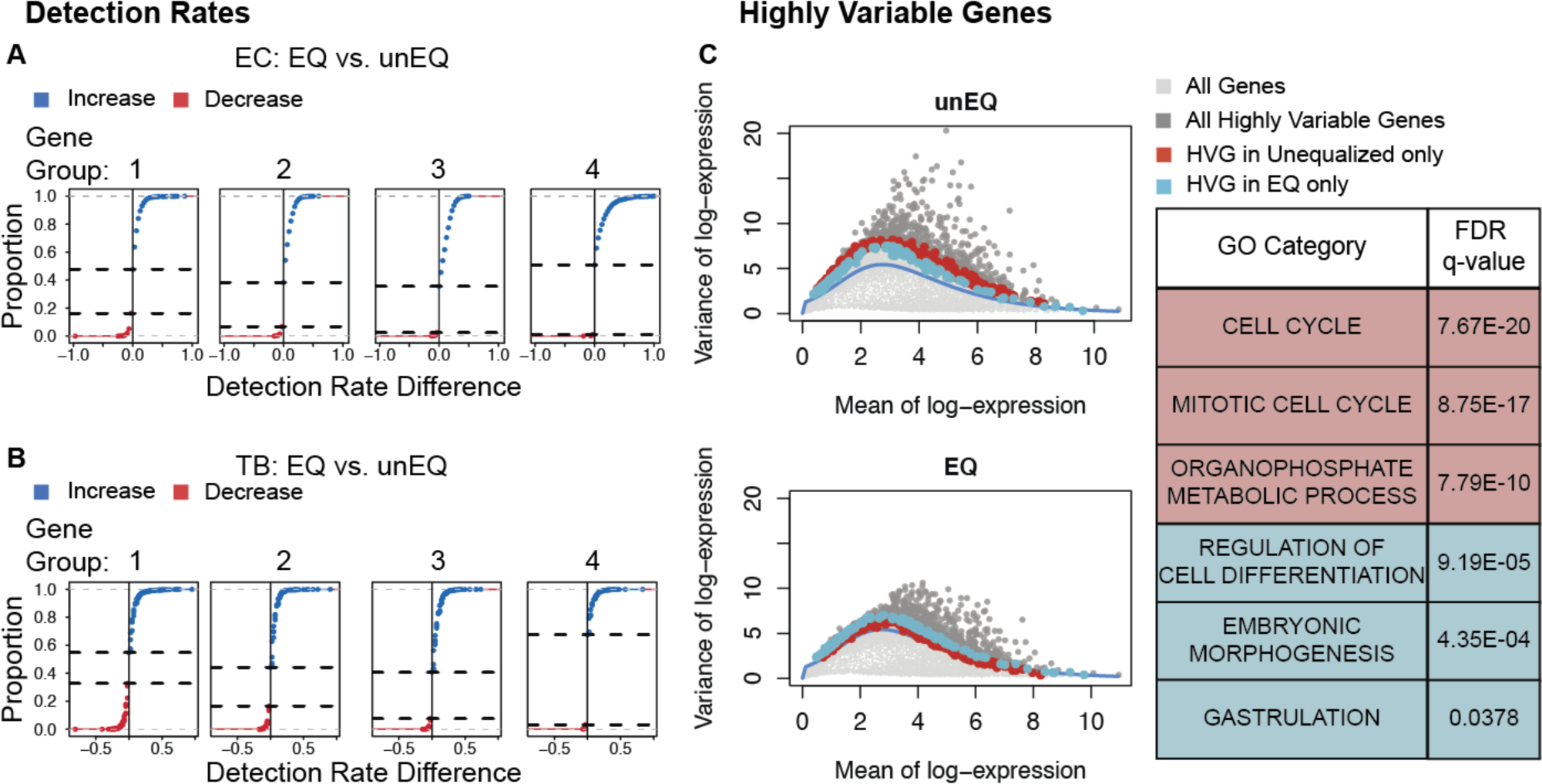
Equalization improves detection rates and decreases expression variability. A.) For the EC dataset, genes were divided into four equally sized groups based on their median nonzero expression. For each gene, the difference between the detection rate in the EQ versus the unEQ experiments was calculated. The cumulative distribution curve is shown for the detection rate differences for genes in each expression group. The two horizonal dotted lines indicate the proportion of genes that decrease in detection rate (bottom line) and one minus the proportion of genes that increase in detection rate (top line). B.) Same as A for the TB dataset. C.) Scatter plot of every gene’s mean and variance for the unEQ (top) and EQ (bottom) datasets (light gray). The smoothed fit line represents technical variability. The mean and variance were calculated over all cells, both EC and TB. Genes having FDR < .10 in either dataset are shown in dark gray. Shown in red are the highly variable genes with FDR < .1 in the unEQ dataset only, and in blue are the highly variable genes with FDR < .1 in the EQ dataset only. In the table are the top three GO biological processes enriched for genes that are only HVG in the unEQ (red) or EQ (blue) experiments.

**Figure 4.**
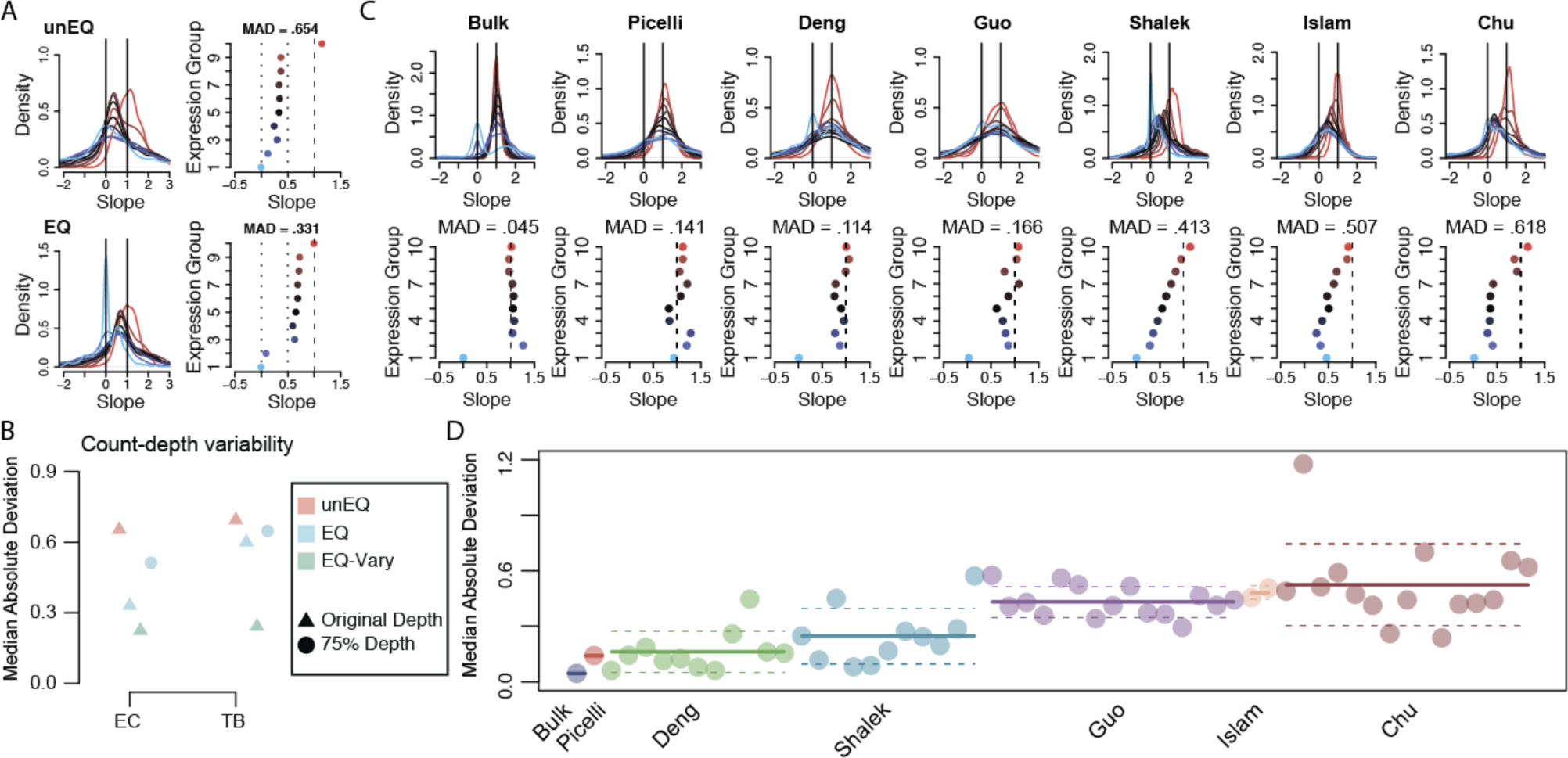
Count-depth rate in equalized scRNA-seq experiments. A.) For the unEQ and EQ EC datasets, the count-depth rate was calculated for all genes as the slope of a median quantile regression. Genes were divided into ten equally sized groups based on their median nonzero expression across all cells in the dataset. B.) The median absolute deviation (MAD) of all experiments slope modes is shown. C.) Same as A for seven representative datasets from seven published studies. D.) Similar to B for all datasets in the seven published studies. The solid line indicates the mean MAD and the dashed line indicates the one standard deviation.

### Simulation Framework

Let *M*_*g,j*_ be the true number of mRNA’s present for gene *g* in cell *j* and has distribution, *M*_*g,j*_*∼Poisson*(*μ*_*g*_), where *g*= 1, …, *G, j*= 1, …, *N*, and *μ*_*g*_ is the true gene-specific expression mean. For scRNA-seq the cell is first isolated, then the mRNA is captured following cell lysis. A reverse transcription step occurs immediately after and converts the mRNA to cDNA. It is currently not possible to naturally estimate these two steps separately. Thus, here we model both of these events together as a single process. The number of molecules successfully captured for genes in cell *j* is represented as:

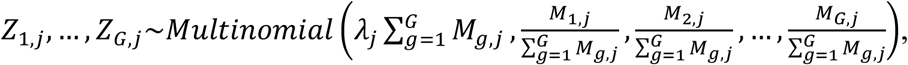

where *λ*_*j*_ is the efficiency of conversion, referred to as the capture efficiency. Following this step, the cDNA molecules are exponentially amplified using PCR. The number of successfully amplified cDNA molecules for gene *g* in cell *j* is: *A*_*g,j*_=*Z*_*g,j*_(1+ *ρ*_*j*_)^*C*^, where C is the number of amplification cycles and *ρ*_*j*_ is the efficiency. If *ρ*_*j*_=1, then all molecules double each cycle. We expect *ρ*_*j*_ to vary across reactions and is independent across cells.

All the following steps occurred in the C1 Fluidigm platform. The next steps involve re-plating the cells for further library preparation. Typically, the cDNA would be quantified to make sure the quality is high. An optional step is to equalize the cDNA concentrations to make them as similar as possible. This is first done by first estimating a small acceptable range of concentrations from the smallest among the cells. One may dilute all concentrations to the smallest observed, or alternatively ensure that the concentrations are within a small range. The median of the range is then the target concentration from which a dilution factor is estimated for all cells outside the range. The dilution factor is estimated as *S*_*j*_*∼Normal*(*τ*_*j*_,*τ*_*j*_*0.1), where *τ*_*j*_ is the target and estimated as:

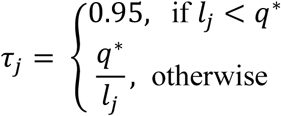

where *l*_*j*_ is the cDNA concentration for cell *j*, and *q*^*^ is the median of the acceptable concentration range. The number of cDNA molecules in cell *j* after equalizing cDNA concentrations is represented here as:

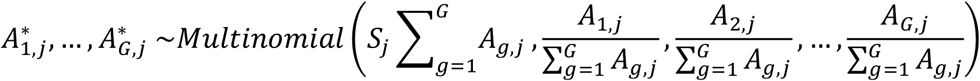

Following the protocols for C1 Fluidigm (Smart-seq and Smart-seq2), next the cDNA is fragmented into shorter pieces and sequencing adapters and cell-specific indexes are added. We model this similar to capture efficiency since the failure of any particular cDNA removes it from further consideration in sequencing. This is commonly referred to as ‘tagmentation’. We denote the tagmentation efficiency here as *γ*_*j*_. The number of cDNA molecules successfully tagmented for genes in cell j is represented as:

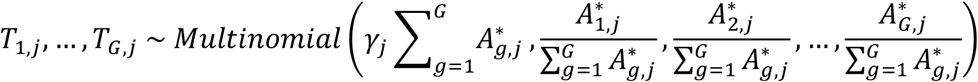

Next, the cDNA molecules go through a second round of PCR amplification, where for gene *g* in cell *j* the number of amplified molecules is represented as:

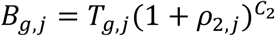, where *C*_2_ is the number of amplification cycles and *ρ*_2,*j*_ is the efficiency per cell. Finally, the observed gene counts per cell, *Y*_*g,j*_, are obtained by:

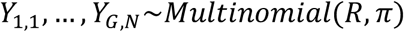

where 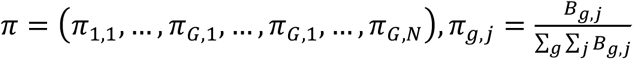, and *R* is the total number of sequences obtained.

### Estimation of simulation parameters

For the simulation framework described above, a number of parameters must be set or estimated. The number of genes and cells were set to match that of the unEQ EC dataset. To estimate the initial gene means we scaled all cells to have a total of 500,000 counts, then estimated the mean for each gene. Since the majority of zeros are thought to occur during the capture step (cell lysis and reverse transcription), the capture efficiency has the largest impact on the detection rates. We estimated the cell-specific capture efficiency for each cell from a Normal distribution with mean 0.078 and standard deviation 0.02. To estimate the mean capture efficiency, we first considered the detection rate per cell as the probability of observing a nonzero, and estimated the average detection rate as one minus the average probability of a gene being zero in the simulated data. Then, for any gene, the probability of a gene being zero was estimated using the Binomial distribution with the probability of detection equal to the ratio of the gene’s mean to the total number of genes in the data and number of trials was set to the expected number of detected genes for a given capture efficiency. Using the optimize function in R, the optimal capture efficiency was that which minimized the distance between the mean probability of detection between the simulated and the unEQ EC data. The standard deviation for the capture efficiency was set to the standard deviation of cell-specific detection rates in the unEQ EC dataset. The first PCR amplification was set to have efficiency from Normal(0.9, .02) with 18 cycles. The equalization for the Unequalized EC dataset was done such that libraries with large cDNA concentrations were diluted to reduce the total range. In the simulation, any libraries with total cDNA counts larger than the 80^th^ quantile among all cells were subsampled based on library-specific dilution factors. Each cell’s target dilution factor was the ratio of its total cDNA to the median of the target range. The cell-specific factors were then estimated from a Normal distribution with the mean being the target dilution factor and standard deviation being 10% of the target dilution factor. The tagmentation step efficiencies were sampled from a Uniform(0.95, 1). The second PCR amplification was set to have efficiency from Normal(0.90, 0.2) with 12 cycles. The total sequencing depth was set to the total counts in the EC dataset.

## Results

### In silico investigation of cDNA equalization using Scaffold

As detailed in Methods and Figure 1A, Scaffold allows for assessment of how each step of the single-cell protocol (cell lysis, amplification, equalization, library preparation, and sequencing depth) affects scRNA-seq measurements. Using an scRNA-seq dataset of unequalized endothelial cells (unEQ EC) as a reference, Scaffold estimated starting parameters and simulated data that reproduced the features of the unEQ EC dataset including gene-specific means, variances, and proportions of zeros (Figure 1B-E). Systematic variability in the count-depth rate, a feature shown to be unique to scRNA-seq data(3), was also reproduced (Figure 1F and Supplementary Figure 1).

Holding all other parameters constant, we used Scaffold to simulate data while varying parameters for equalization and sequencing depth and found that cDNA equalization has the largest effect on the average variability in the count-depth rate (Supplementary Figure 1C&D), while the total sequencing depth (Supplementary Figure 1E) had little effect.

To examine the effect of equalization on other properties of the data, we simulated additional datasets with and without equalization holding all other steps constant. Specifically, we simulated pairs of unequalized and equalized datasets by adjusting only the equalization parameter. In simulated datasets, gene-specific variation decreased by an average of 16.5% due to equalization alone and the variability in the sequencing depths was reduced by 60.9% despite the simulations having the same average depth (Figure 1G).

### Experiments to assess the effect of cDNA equalization

Given results from the simulation study, we hypothesized that a lack of equalization during the preparation of single-cell libraries would increase variation in the amount of input cDNA which in turn could contribute to reduced gene detection and increased variability in expression estimates observed in scRNA-seq data. To test this hypothesis, we applied alternative protocols to full-length single-cell cDNA libraries of identical cells to generate matched scRNA-seq data sets (Fig. 2A). The original data includes single endothelial cells (EC) and trophoblast-like cells (TB) derived from human embryonic stem cells (hESC)(22) which were unequalized (unEQ). For these experiments, the cDNA input ranged from 0.125 - 0.375 ng (Methods). In the next series of experiments, we equalized the same set of single-cell cDNA to a fixed input (0.1 ng) across all the cells. Prior to sequencing, cells were pooled at an equal volume (EQ) or pooled by a scaling factor to produce highly variable sequencing depths (EQ-Vary) (Figure 2A). Finally, we replicated the entire EQ experiment, including equalized cDNA input and pooling, but we sequenced at approximately three-quarters the depth of the previous experiments (EQ-75%).Because these four conditions all derive from identical cells, these experiments provide the most robust investigation to date on how input cDNA variations impact scRNA-seq data.

### Equalization increases cell-specific and gene-specific detection rates

A common challenge in scRNA-seq experiments is the high proportions of zeros, or dropouts. Dropouts are due to an incomplete sampling process, stochastic gene expression, and inefficient capture of mRNA, with the probability of dropping out inversely related to a gene’s underlying expression level(23). Equalizing cDNA libraries would not recover dropouts that occur upstream in a protocol, but it may recover dropouts that are due to inefficiencies in later preparation steps (e.g. second PCR amplification) or due to underrepresentation in the pooled library. Thus, we first investigated the effect of cDNA equalization on cell-specific detection rates, defined as the proportion of nonzero genes within a cell. Across both EC and TB cells, we observed an increase in the efficiency of gene detection in the equalized experiments (Figure 2B). An average of 745 (8.6%) more genes per cell were detected with expression greater than zero in the EQ versus the unEQ experiments. EQ-vary, which was pooled in a way to reflect possible inefficiencies that might occur after equalization such as during pooling or amplification, reduced the detection efficiency slightly to 534 (6.2%) more genes detected on average. Comparatively, the effect of equalization on gene detection is stronger than the effect of solely increasing total sequencing depth. Between EQ and EQ-75%, in which both experiments were equalized but the latter had three-quarters the sequencing depth, we observed 470 (5.0%) fewer genes detected per cell in EQ-75%.

We further investigated the gene-level detection rate across experiments, defined as the proportion of cells with nonzero expression for each gene (Figure 3A&B). Here we calculated the difference in gene-level detection rates between EQ and unEQ while accounting for differences in sequencing depth (Methods). The overall increase in detection efficiency due to equalization translates to a 31.1% increase in genes having consistent detection in all EC cells and a 17.9% increase in TB cells (1002 and 622 genes, respectively). We also observed a 10.4% decrease in the number of genes not detected in any cells for EC and an 8.1% decrease in TB (382 and 276 genes, respectively).

Since a gene’s detection rate is related to its expression level, we further analyzed detection differences by splitting genes into four equally sized gene groups based on their nonzero median expression. We first assessed what differences would appear due to random chance by randomly splitting the EC or TB cells in the unEQ dataset into two groups and examined the detection rate differences between them. We observed approximately equal proportions of genes having increased/decreased detection rates across all expression groups for both experimental conditions (Supplementary Figure 2).

Between the EQ data and unEQ datasets, we consistently see a higher proportion of genes having a higher detection rate in the equalized dataset especially among the moderately expressed genes (62% and 64% for EC gene groups 2 and 3; 56% and 59% for TB gene groups 2 and 3)(Figure 3A&B). The average increase in detection rate in the equalized experiments for the genes in Groups 2-4 is 13.6% in EC2 and 7.9% for TB2. In comparison, we performed the same analysis between the EQ and EQ-Vary datasets which underwent the same equalization procedure and found the ratio of genes with increasing versus decreasing detection rate was stable across expression groups; the increase variability in sequencing depth did not compromise the detection rate in the equalized dataset (Supplementary Figure 3).

To identify any functional relevance of genes with increased or decreased detection rates in the EQ experiment we performed gene-set enrichment using MSigDB’s list of GO biological processes on the top 200 genes sorted by their magnitude change in detection. Genes with increased detection rate in the EQ experiment were enriched for important developmental processes including morphogenesis, and tube and epithelium development in both EC and TB (Supplementary Table 1). Genes with decreased detection rates after equalization tended to be among the most lowly expressed genes. Of the 200 genes with the most decreased detection, 142 were in the lowest expression group in EC and 162 such genes in TB. Taken together, these results suggest that equalization improves the detection of biologically relevant genes without compromising signal.

### Equalization reduces nuisance variation

Next, we investigated the effect of equalization on gene expression variability. A common first step in single-cell clustering or trajectory inference analysis is to reduce the data to the most informative set of genes often defined as the most highly variable genes (HVG). However, in the presence of excess nuisance variation, the top ranked HVG may not reflect the most relevant set of genes. Here, we detected HVG by decomposing the total variance of each gene into technical and biological components. To do so, we estimated a mean-dependent trend for the mean-variance relationship across all genes to represent technical variability (Methods). A gene’s biological variability was calculated as the distance between a gene’s total variability and its fitted trend value. An HVG classification was assigned to genes having biological variability significantly larger than zero (FDR < .10). HVG genes in the unequalized experiment were enriched in GO biological processes involving the cell cycle. This is likely due to the fact that cellular mRNA content is directly related to cell cycle stage and, consequently, if cDNA content is not equalized across cells, variability in cell cycle genes is prominent in the resulting data. Following equalization, genes classified as HVG were enriched for biological processes specific to EC cells including gastrulation and cell fate/differentiation (Figure 3C & Supplementary Table 2).

### Equalization reduces technical artifacts in the count-depth rate

Previously, we reported that scRNA-seq data display systematic variation in the relationship between a gene’s observed expression and sequencing depth (which we termed the count-depth rate), whereby a gene’s expected increase in expression with increased sequencing depth fails to materialize(3). Variability in the count-depth rate affects downstream analysis as popular scale-factor based normalization methods assume that the count-depth rate is common across genes and equal to one on the log-log scale (3,24).

As shown in Bacher et al., 2017, much of the variability in the count-depth rate arises from under-detection of genes despite increasing sequencing depth since highly expressed genes are over-represented during sequencing. Since equalizing cDNA increases detection rates, we hypothesized that it may also reduce variability in the count-depth rate. To investigate, we quantified the count-depth rate for every gene using median quantile regression, where a slope of one indicates a proportional increase of gene expression with sequencing depth (Supplementary Figure 4). Next, we binned genes into ten equally sized groups based on their median nonzero expression. In the unEQ dataset, we found only highly expressed genes had slopes near one and slopes gradually decreased with gene expression level (Figure 4A). The extent of variability in the count-depth rate was measured using the median of the absolute deviations MAD) of the ten groups slope mode from their expected value of one. The EQ experiments had a lower MAD and displayed less variability in the count-depth rates for both EC and TB (Figure 4A&B). EQ-75% was similar to the EQ datasets, indicating the count-depth rate is not affected by total sequencing depth. The EQ-Vary experiment had the most reduction in count-depth variability, with most slopes close to 1 (Supplemental Figure 5), due to its increased dissociation of cell size with sequencing depth.

As more single-cell datasets have become public and identically processed in databases such as conquer(25), we were able to inquire whether systematic variability in the count-depth rate was reduced across scRNA-seq data in published studies. Across seven different studies, we found large heterogeneity in the experiment-specific count-depth rates with the MAD ranging from 0.045 to 1.176 (Figure 4C&D). We found no revealing association between the average MAD within study and various properties of the scRNA-seq data, including the average sequencing depth, cell-specific detection rate, organism, or number of cells (Table 1). However, consistent with our simulated and experimental datasets, the publicly available studies in which equalization was performed had significantly lower MAD values (p-value < .001), higher cell-specific detection rates (p-value < .001), and higher gene-specific detection rates (p-value = .039) (Supplementary Figure 6). On average the equalized datasets contain 2,215 additional genes detected consistently in every cell compared to the unequalized datasets (p-value < .001 & Supplementary Figure 6).

## Discussion

Obtaining the highest quality data with minimal technical variability remains a goal for scRNA-seq experiments. Given the competitive nature of the sequencing process, transcripts that are highly expressed are often overrepresented in the final library and will consume a large proportion of the total reads leading to low detection rates for the majority of genes. Here we showed that equalizing single-cell cDNA libraries prior to pooling decreases nuisance variation such as that attributable to cell cycle while improving the detection rate and reducing variability in biologically relevant genes.

Our finding of reduced variability in expression for cell cycle genes in equalized experiments is novel, yet not unexpected since cell cycle signals are often the largest drivers of differences in total mRNA. Note that if cell cycle signals are of marked interest then equalization may not be appropriate. However, reduction of cell-cycle signals has been implemented in most scRNA-seq analysis pipelines as it is considered a hindrance in downstream analysis(26,27).

In many cases, identified sources of technical variability in downstream analyses have proven to be excellent targets for protocol improvement(28–31). Scaffold, our simulation framework, offers an opportunity to directly and efficiently explore how different steps in a protocol affect scRNA-seq data. Here, we focused the effect of equalizing cDNA concentration across cells. However, Scaffold provides a framework to study other parameters, or to simulate data that recapitulates characteristics of scRNA-seq data (e.g. detection rates and count-depth rate).

In practice, the process of equalizing cDNA concentrations is non-trivial and time-consuming, leading it to be one of the critical limiting points of the library preparation process(32). Automation has alleviated this to some extent, and has been used in large single-cell sequencing projects such as the Tabula Muris(33). However, some state-of-the-art protocols, such as 10X, profile scRNA-seq measurements from thousands to millions of cells using massively parallel sequencing systems with high levels of multiplexing (Lundin et al. 2010) and equalization is not possible since cDNA is pooled early in the experiment. We expect that single-cell protocols will continue to advance and improve with technology. Our study offers insight into one mechanism worth further exploration in protocol design and development.

## Data availability

All R code used for analysis or simulations is available at https://github.com/rhondabacher/scEqualization-Paper. The simulation package Scaffold is available at https://github.com/rhondabacher/scaffold. The unEQ, EQ, EQ-Vary, and EQ-75% datasets are available at the NCBI Gene Expression Omnibus: GSE156494 (https://www.ncbi.nlm.nih.gov/geo/query/acc.cgi?acc=GSE156494; the reviewer access token is ehipcsoybvgtrud).

For the publicly available datasets, we obtained processed counts from the conquer scRNA-seq database for three single-cell RNA-seq datasets processed identically: Deng et al., 2014 (38), Guo et al., 2015 (39), and Shalek et al., 2014 (40). The Chu et al., 2016 (22) data was obtained from the Gene Expression Omnibus (GEO) with the accession number GSE75748. The Islam et al., 2011(41) data was obtained from GEO with the accession number GSE29087. The H1-bulk data from Bacher et al., 2017 (3) was obtained from GEO with the accession number GSE85917. The Picelli et al., 2013 (42) was obtained from the GEO with the accession number GSE49321.

## Acknowledgements

We thank J. Steill and S. Swanson for initial RNA-seq read processing.

## Author contributions

R.B. and C.K. conceived and designed the research and wrote the manuscript. L.-F.C. and J.B. conceived, designed and performed experiments. R.B. processed and analyzed all datasets. P.K. contributed to simulation code development. R.S. and J.A.T were involved in planning and supervising experiments. All co-authors contributed to the writing of the manuscript.

