## Supplemental Figures for "Enhancing biological signals and detection rates in single-cell RNA-seq experiments with cDNA library equalization"

Supplementary Figure 1. A.) Density plots of the distribution of estimated count-depth rates for the unEQ EC dataset with genes grouped by expression level (left) and the mode of each group's slope distribution (right). The median absolute deviation (MAD) of the slope modes from one is used to quantify the variability in the count-depth rate. B.) Similar to A for one simulated dataset with parameters set to match the unEQ EC dataset. C.) Same as B but the simulation was done assuming equalization of cDNA concentrations. D.) The MAD is shown for 100 simulations holding all parameters constant, and varying the amount of equalization. E.) The MAD for 100 simulations holding all parameters constant, and varying the sequencing depth as X times the original, where X = 0.25, 0.5, 1, 2, 3 .

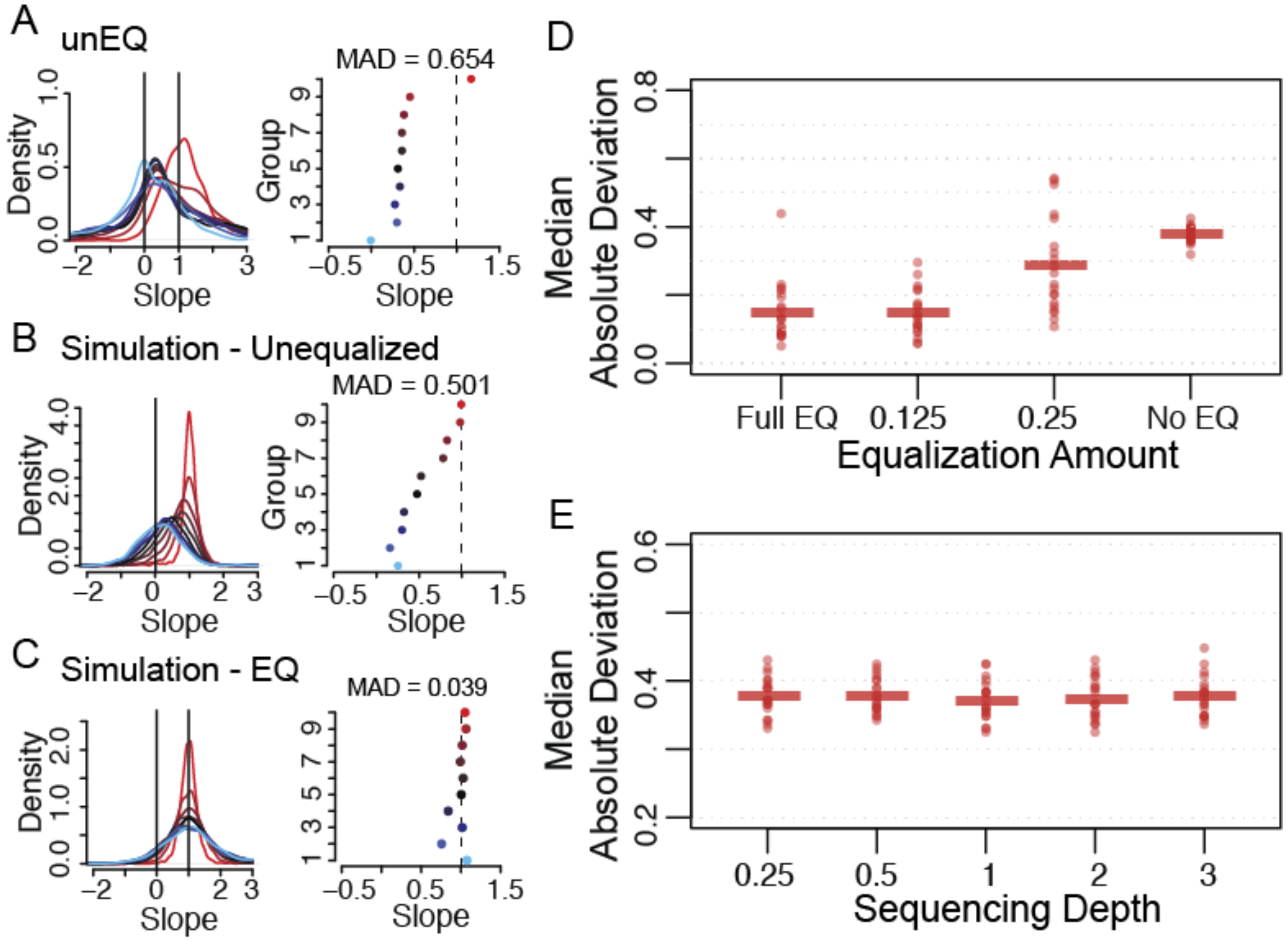

Supplementary Figure 2. Detection rate differences for a random split of cells in each of the EC and TB unEQ datasets. Genes were divided into four equally sized groups based on their median nonzero expression. For each gene, the difference between the detection rate in the random data splits was calculated. The cumulative distribution curve is shown for the detection rate differences for genes in each expression group. The two horizonal dotted lines indicate the proportion of genes that decrease in detection rate (bottom line) and one minus the proportion of genes that increase in detection rate (top line).

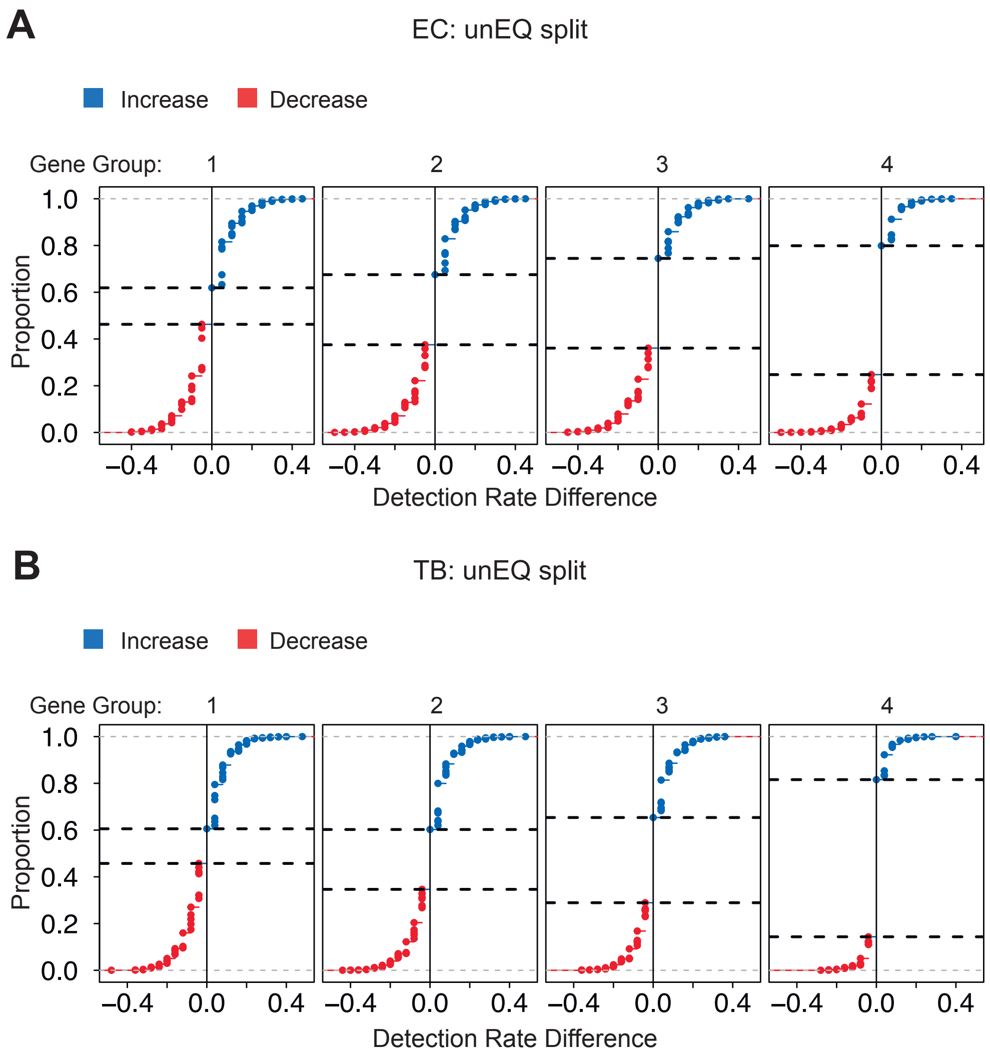

Supplementary Figure 3. Detection rate differences between the EQ and EQ-Vary datasets for the EC and TB conditions. Genes were divided into four equally sized groups based on their median nonzero expression. For each gene, the difference between the detection rate in the EQ versus EQ-Vary experiments was calculated. The cumulative distribution curve is shown for the detection rate differences for genes in each expression group. The two horizonal dotted lines indicate the proportion of genes that decrease in detection rate (bottom line) and one minus the proportion of genes that increase in detection rate (top line).

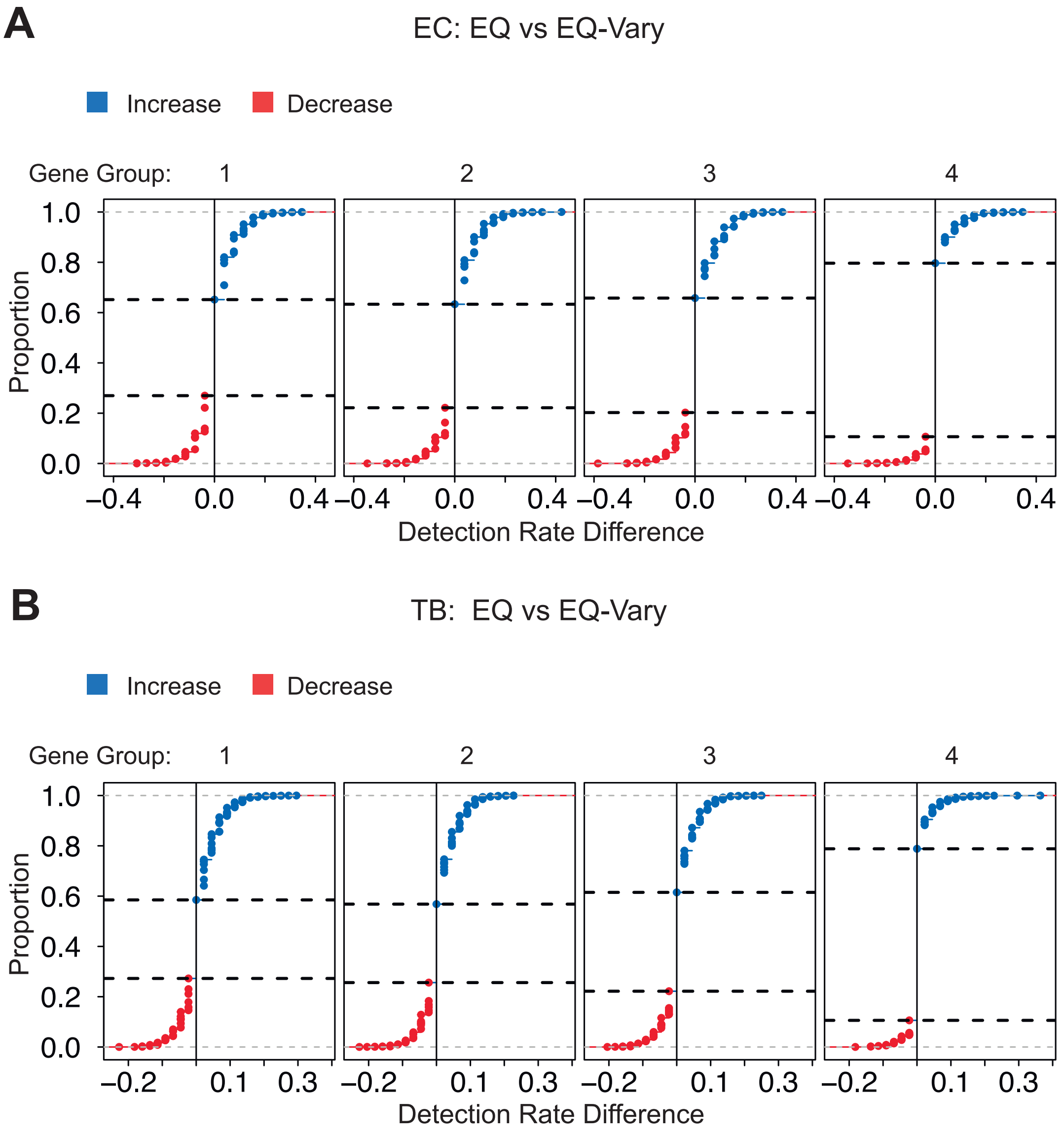

Supplementary Figure 4. The count-depth rate is estimated as a median quantile regression of log expression versus log sequencing depth. A low, moderate, and highly expressed gene are shown having a count-depth rate of 0.05, 0.53, and 0.96, respectively.

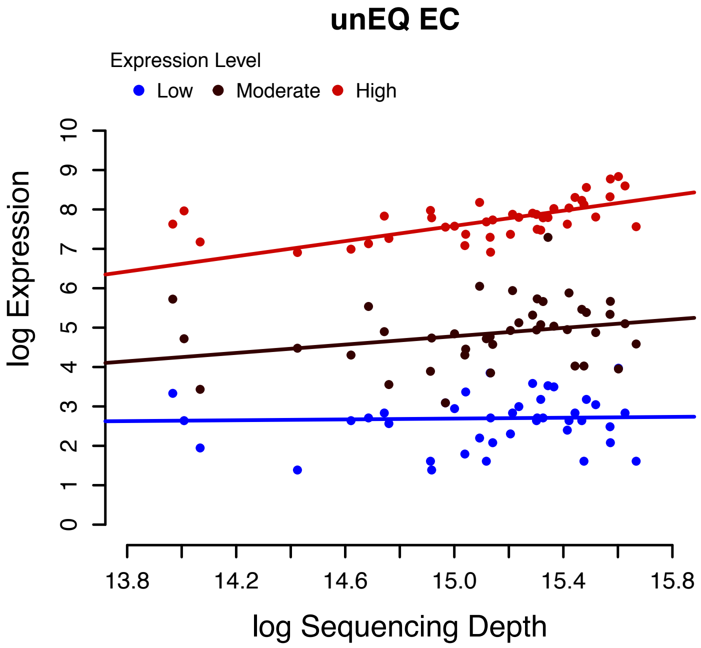

Supplementary Figure 5. Count-depth relationships for all EC and TB experiments. A) Density plots of the distribution of estimated count-depth rates for the EC datasets with genes grouped by expression level (left) and the mode of each group's slope distribution (right). B) Similar to A for the TB datasets.

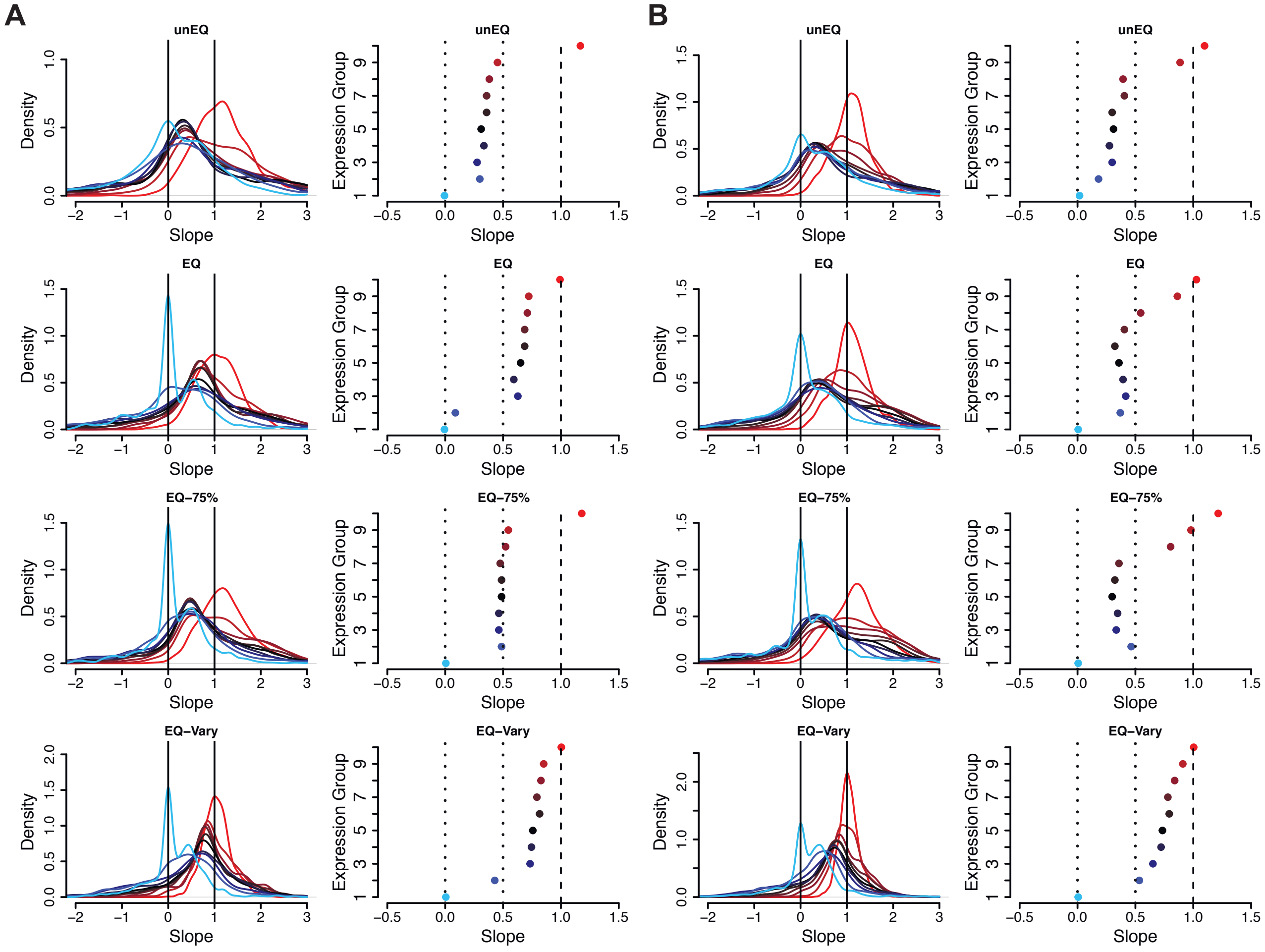

Supplementary Figure 6. Cell-specific and gene-specific properties are shown for pairs of equalized and unequalized datasets. Pairs of unequalized experiments were also simulated and compared to demonstrate the percent of change due to random sampling.

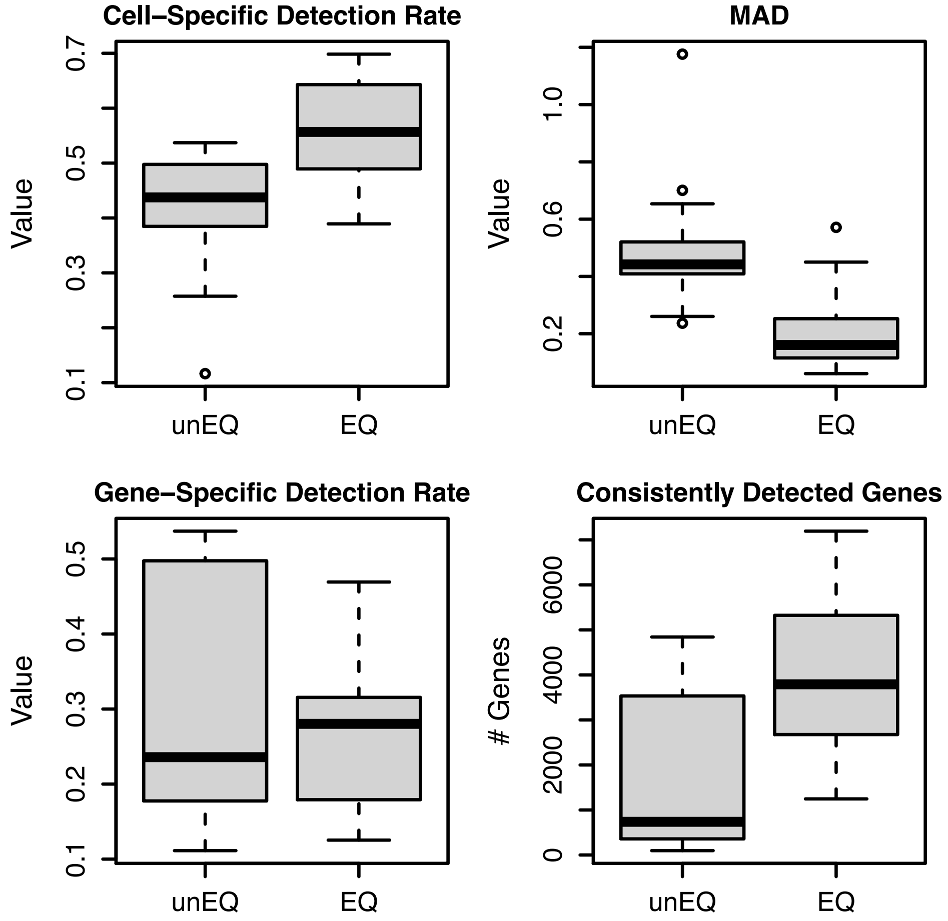

Supplementary Figure 7. Cell-cell Pearson correlations across experiments. Cells are in the same order along the axes for the two experiments being compared. The strong diagonal correlation verifies that the cells are indeed the same across experiments.

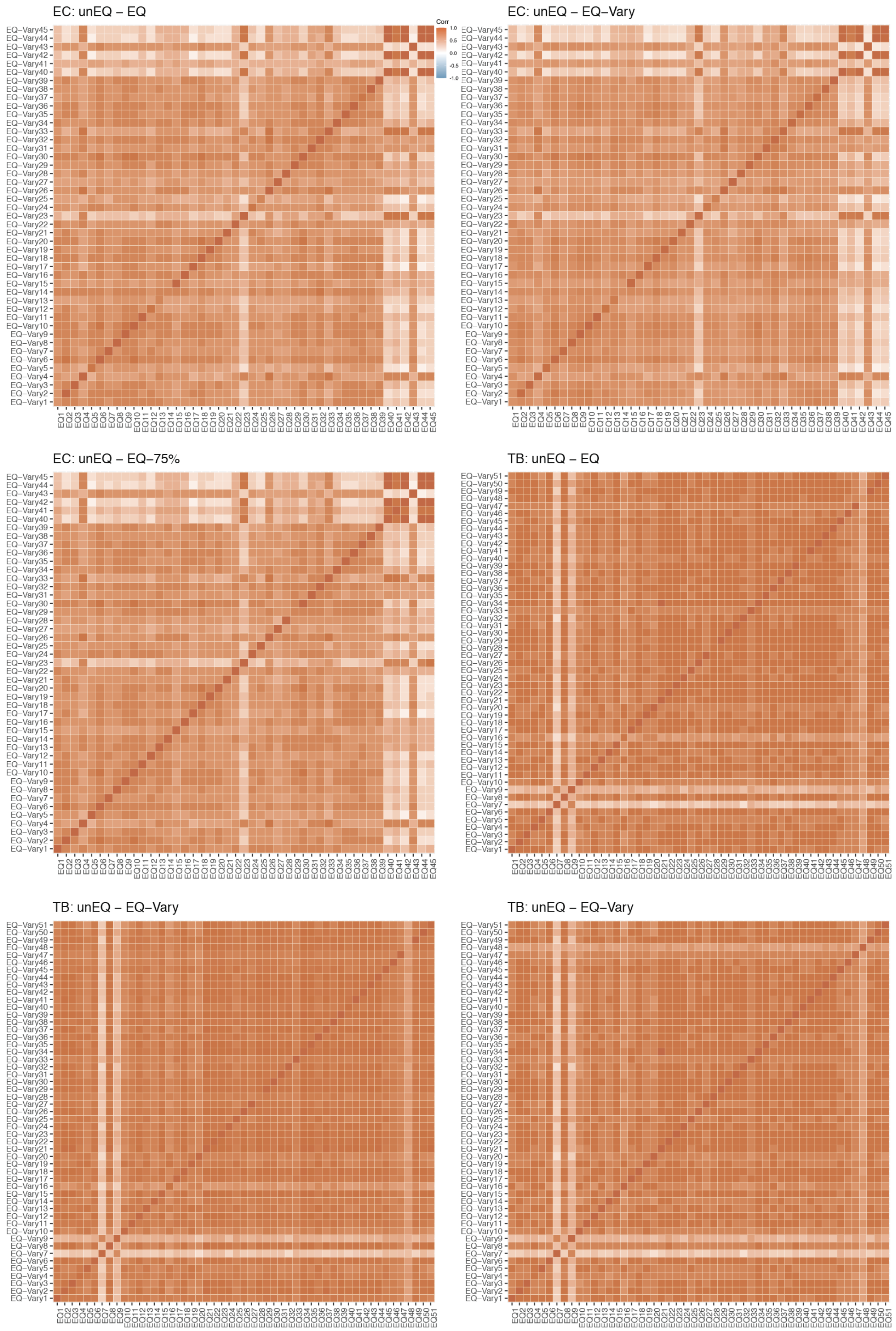

Supplementary Figure 8. Outliers cells were removed from further analysis. Outliers were identified as those having log10 total counts < 5.4 or the percent of counts in the top 50 genes was > 31%.

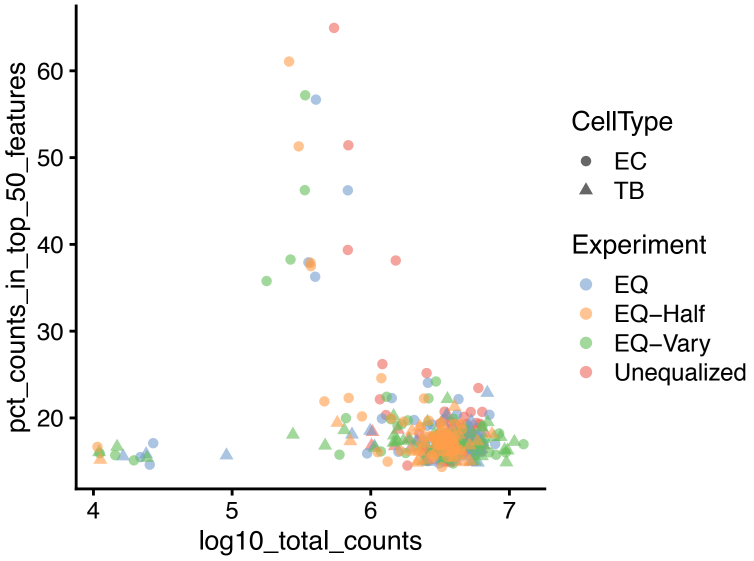
